# Beyond Temporary Scaffolding: Persistent Hippocampal Engagement and Brain-Wide Network Reorganization in Remote Memory Consolidation

**DOI:** 10.64898/2026.07.26.740756

**Authors:** Ruimin Wang, Ruedeerat Keerativittayayut, Shuya Morioka, Masaki Takeda, Isao Hasegawa, Koji Jimura, Kiyoshi Nakahara

## Abstract

The standard model of memory consolidation posits a temporary role for the hippocampus. We challenge this view by combining a novel face–name associative memory task with cross-modal decoding of fMRI data. We demonstrate that the hippocampus remains persistently engaged in representing both recent and remote memories, with distinct subregional contributions: recent memories were represented in the right posterior hippocampus, whereas remote memories additionally recruited the left anterior hippocampus. This sustained hippocampal involvement occurred alongside a large-scale network reorganization. Remote memory retrieval was accompanied by increased global connectivity among semantic regions, yet the hippocampus itself did not emerge as a major hub. Instead, network hubs shifted from the medial prefrontal cortex (mPFC) for recent memories to the mPFC and bilateral inferior parietal lobule for remote memories. These findings reveal a dynamic process in which the hippocampus preserves long-lasting representational functions while the broader memory network undergoes extensive reorganization.

## Introduction

Our sense of self is grounded in long-term declarative memories—personal experiences (episodic memory) and accumulated knowledge (semantic memory). The hippocampus (HPC) is essential for encoding these memories and is thought to guide their gradual reorganization across the neocortex through a process known as systems consolidation ^1, 2^.

Evidence for systems consolidation comes from cases of temporally graded retrograde amnesia (RA), where hippocampal damage makes recent memories more vulnerable than older ones ^1, 3^. Yet, clinical findings are inconsistent ^4^. For instance, patient HM exhibited severe damage to memories from recent years following the removal of the medial temporal lobe (MTL), including the HPC ^5^. In contrast, patient VC, with more selective hippocampal damage, was unable to recall even childhood episodes ^6^.

This remarkable variation has fueled a long-standing debate in neuroscience, leading to two main theories of systems consolidation: the standard consolidation theory (SCT) ^7^ and the multiple trace theory (MTT) ^8, 9^. SCT posits that memories are temporarily retained in the HPC before being transferred to the neocortex for permanent storage ^7, 10^, transitioning from episodic to semantic memory ^11^. In contrast, MTT argues that while semantic memories can become independent of the hippocampus, detailed episodic memories remain dependent on the HPC indefinitely ^12^. Trace Transformation Theory (TTT) refines this idea, suggesting that the posterior hippocampus (pHPC) supports detailed memory retrieval by connecting to the posterior neocortex, while the anterior hippocampus (aHPC) is linked to the anterior neocortex for gist-based memory retrieval ^2, 13^. Furthermore, TTT emphasizes the increasing importance of schema—structured frameworks of prior knowledge—in the retrieval of episodic memory over time, where the ventromedial prefrontal cortex (vmPFC) plays a central role in the memory retrieval network ^14^. However, empirical findings remain mixed. For instance, Sommer reported that vmPFC involvement diminished after months of repetition-based learning, while semantic regions such as the left inferior frontal gyrus (IFG) became increasingly engaged ^15^. These inconsistencies highlight three unresolved questions in systems consolidation: 1. Does the hippocampus retain representational roles for remote memories?; 2. How does large-scale network organization change between recent and remote memory retrieval?; and 3. Which brain regions act as central hubs in these reorganized networks?

Many previous neuroimaging studies have relied on famous face recognition and autobiographical interviews to assess remote memories from years ago. While some studies using univariate analysis support the SCT by showing decreased HPC activation for remote memories ^16, 17, 18^, others support the MTT/TTT by finding no significant difference ^19, 20, 21, 22^. These inconsistencies may stem from two issues: the highly individualized nature of autobiographical memories ^18, 19^ and the limits of univariate approaches, which capture overall activation rather than representational content ^23^.

Multivariate pattern analysis (MVPA) provides direct insight into neural representations of individual memories, potentially addressing discrepancies seen in traditional univariate analyses. ^23, 24, 25, 26^. Several recent studies have employed MVPA to examine the role of the HPC in maintaining remote memories by decoding them from fMRI data during memory retrieval ^21, 22^. In these studies, the same memories were recalled multiple times, with one subset used for training a classifier and another subset used for evaluating the classifier’s performance. This approach is commonly known as cross-validation classification. However, due to the highly individualized nature of autobiographical memories, classification techniques based on cross-validation cannot specify which components of memory are accurately classified ^23^. Moreover, HPC activity during memory retrieval is affected by factors such as the repetition of experiences ^15, 17^, the contents being retrieved ^27^, and the type of retrieval cues ^28, 29^. Precise control of memory components is therefore essential ^23^.

To overcome these challenges, we devised a novel face-name associative memory task combined with MVPA and a cross-modal decoding approach. Our task was specifically designed using the faces and names of Pokémon characters, leveraging a unique population: university students with extensive childhood experience of the 2006 Pokémon games (remote memories) but little exposure to the 2019 version (recent memories). Prior work shows that extensive childhood exposure to Pokémon shapes shared cortical representations across adults ^30^, making this an ideal paradigm for controlling both stimulus familiarity and memory age. At the same time, our task also allowed us to perform MVPA with cross-modal (face-name) decoding. Our cross-modal decoding approach further ensured precise representational control beyond conventional cross-validation.

Using this paradigm, we tested whether the hippocampus retains representational traces of remote memories, how the retrieval network reorganizes over time, and which regions emerge as communication hubs. We find that both recent and remote memories are represented within the hippocampus, but through distinct subregions. At the same time, large-scale network organization undergoes striking reconfiguration—from perceptual dominance during recent retrieval to semantic integration during remote retrieval—with hub roles shifting from mPFC alone to mPFC and bilateral inferior parietal lobule. Together, these findings support and refine TTT by revealing persistent hippocampal involvement within a dynamically reorganizing memory network.

## 2. Results

### 2.1 Participants’ experience regarding 2006- and 2019-Pokémon

We analyzed data from 30 participants (mean age: 19.8 ± 1.7 years; six females), collected between December 2019 and September 2020. On average, they began playing the 2006 Pokémon version 11.0 ± 2.0 years ago (between 2006 and 2013) and stopped 6.5 ± 3.5 years ago (between 2008 and 2020) (Fig. 1a). By contrast, participants had limited experience with the 2019 version; 12 had no prior knowledge, while the remaining 18 had at least some exposure to it. On average, participants’ remote memories of the 2006 version were approximately 11.0 ± 2.0 years old, whereas their recent memories of the 2019 version were 0.2 ± 0.3 years old (Fig. 1a).

**Figure 1.**
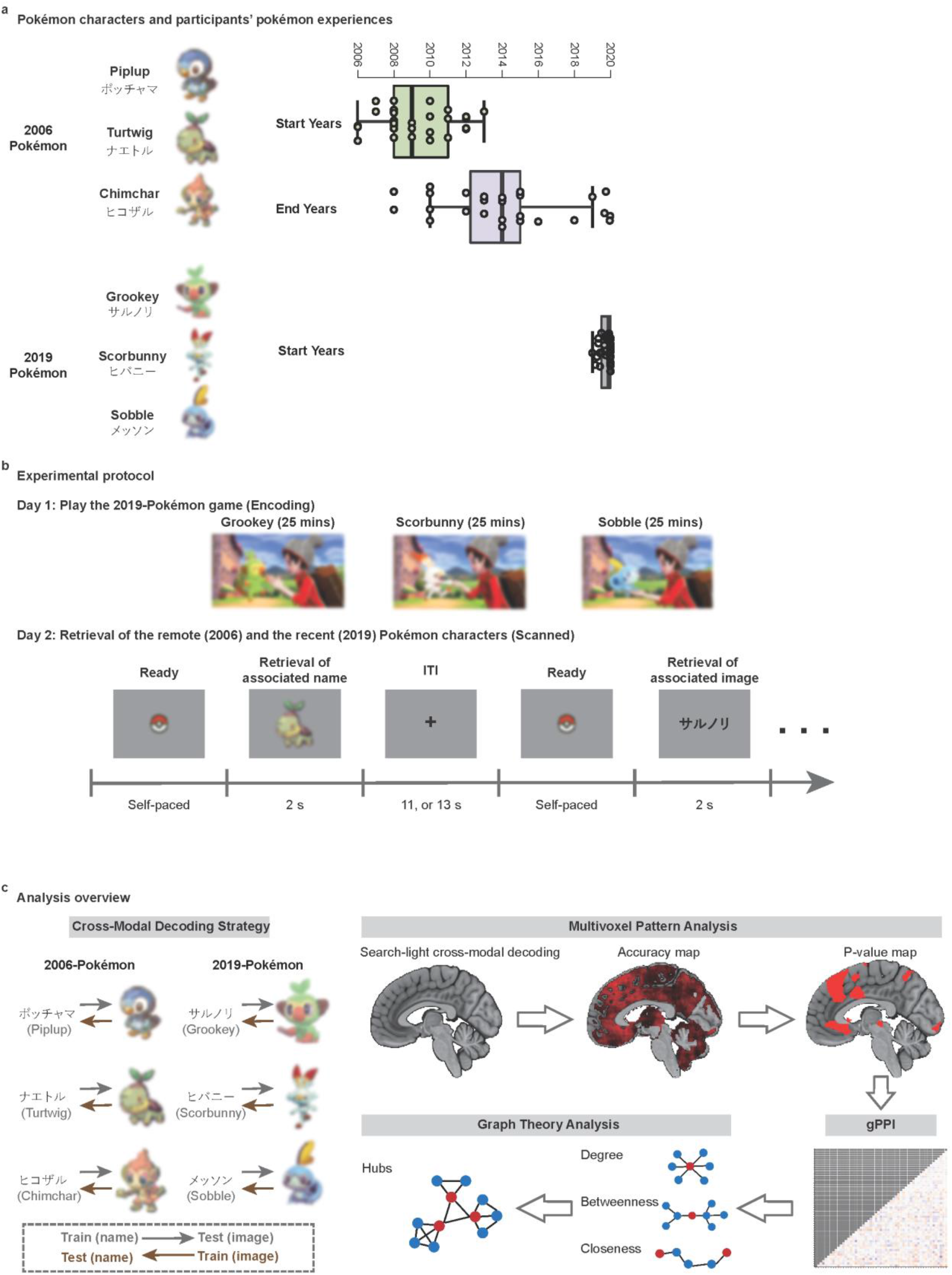
Experimental design and analytic framework. **a**. Three characters each from the 2006 and 2019 Pokémon series were used to probe remote and recent memory, respectively. Box plots summarize participants’ prior experience with Pokémon (*n* = 30 participants; line: median; box: interquartile range (IQR); whiskers: 1.5 × *IQR*) **b**. Experimental protocol. Day 1: Participants interacted with three 2019 Pokémon characters (25 min per character). Day 2: Participants underwent fMRI scans while performing a face–name associative memory task. Each trial began with a Pokéball cue, prompting a right-handed button press to initiate the trial. A face or name of a Pokémon character was then presented for 2 s, and participants retrieved the corresponding associate from memory. Trials were separated by a jittered inter-trial interval (ITI) of 11 or 13 s. **c**. Analytic pipeline. Multivariate pattern analysis (MVPA) was employed with a cross-modal decoding strategy (left) to independently decode remote and recent memories across the whole brain. Brain regions demonstrating decoding accuracy significantly above chance were identified as memory-representation regions. Functional connectivity among these regions was estimated using generalized psychophysiological interaction (gPPI) analysis, performed separately for remote and recent memory retrieval. Finally, graph-theoretical network analysis identified neural hubs supporting remote and recent memory retrieval. For copyright reasons, image related to Pokémon have been blurred.

### 2.2 Memory accuracies of 2006- and 2019-Pokémon characters

The experiment was conducted over two consecutive days (Fig. 1b). On Day 1, participants played the 2019 Pokémon game, where they incidentally learned associations between the faces and names of three selected characters. On Day 2, they performed a face–name association task during fMRI scanning, using the same three characters from both the 2006 and 2019 versions (Fig. 1b). After scanning, participants completed a surprise name-writing test in which they were shown the characters’ faces and asked to recall their names. Participants achieved an average accuracy of 94.4 ± 12.6% for the 2006 characters and 99.0 ± 5.9% for the 2019 characters. There was no significant difference between remote and recent memory retrieval accuracy (paired t-test, *P* = 0.10), indicating comparable precision in retaining Pokémon character names from both versions.

### 2.3 Cross-Modal MVPA Results

We applied a cross-modal (face–name) decoding multivariate pattern analysis (MVPA) using an SVM classifier to identify neural representations of remote and recent associative memories (see Methods 4.5). Specifically, the SVM classifier was trained on brain activation patterns elicited during the retrieval of either the Pokémon names or faces. Its decoding accuracy was then evaluated on the brain activation patterns associated with the other modality of memory items. Brain regions where the decoding of the three 2006 Pokémon characters was significantly above chance level (*P* < 0.01, FDR corrected) were considered to contain neural representations of remote memories. Conversely, regions that successfully decoded the three 2019 Pokémon characters were deemed to maintain neural representations of recent memories

#### 2.3.1 The HPC maintained neural representations of both remote and recent memories

MVPA revealed neural representations of remote associative memories in two hippocampal regions: the left anterior hippocampus (aHPC.L) and the right posterior hippocampus (pHPC.R) (Fig. 2a; Supplementary Table 2). The pHPC.R also contained neural representations of recent memories (Fig. 2b; Supplementary Table 3), with an observed overlap between remote and recent representations in this region (Fig. 2c; Supplementary Table 4). These findings provide evidence that the pHPC sustains memory representations for over a decade and that new representations emerge in the aHPC during systems consolidation. Together, they support key tenets of both the multiple trace theory (MTT) and the trace transformation theory (TTT).

**Figure 2.**
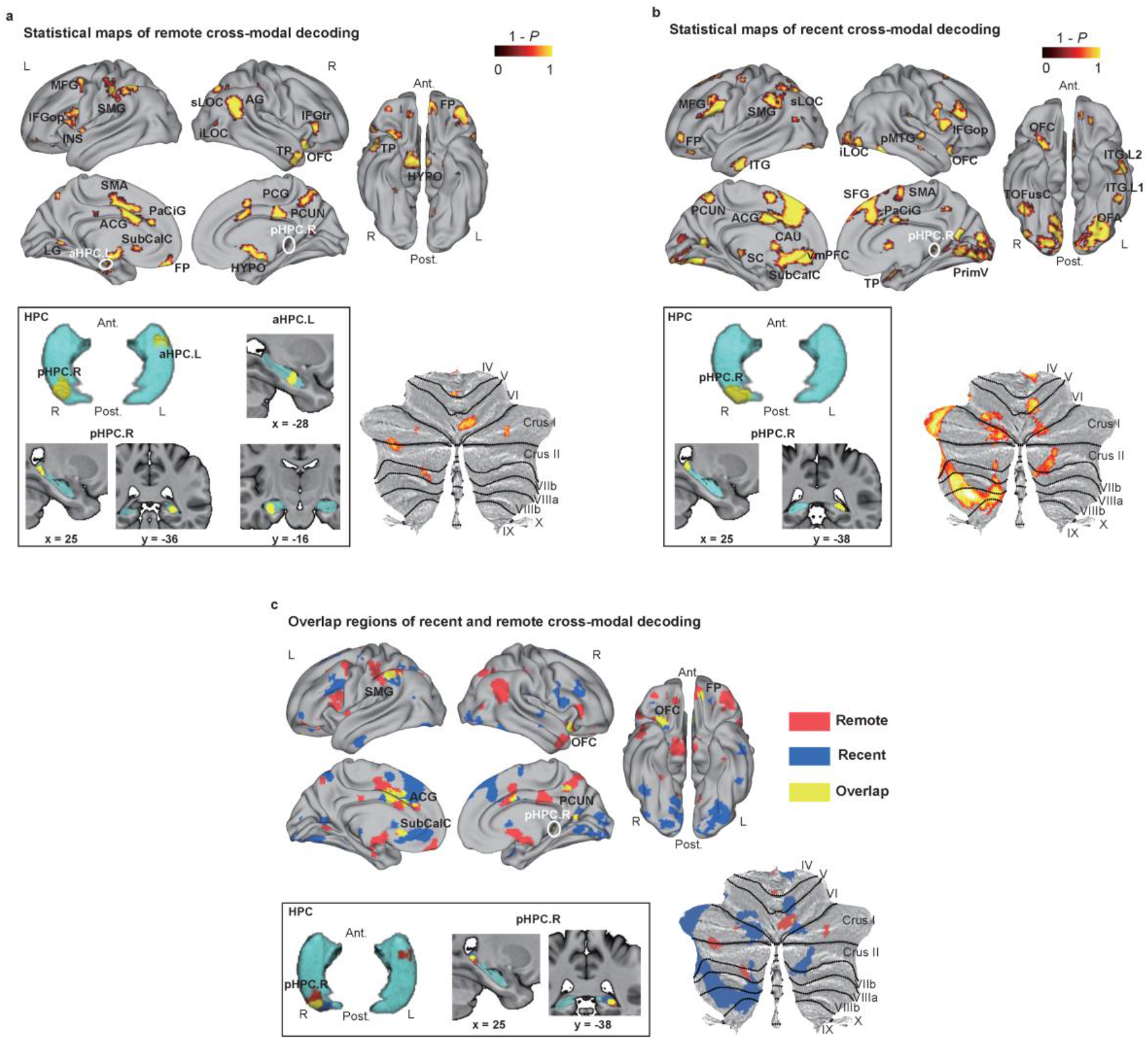
Neural representations of remote and recent memories revealed by cross-modal decoding. **a, b.** Statistical maps (voxel-wise one-sample t-test, 10,000 permutations, *P* < 0.01, FDR-corrected), obtained from cross-modal decoding MVPA, revealed distinct neural representations of remote (**a**) and recent (**b**) memories throughout the brain. Remote memories were predominantly distributed in the frontal-parietal cortices and the limbic system, whereas recent memories exhibited a wide distribution in the frontal and occipital cortices, as well as the cerebellum. **c.** Overlap regions, indicating shared neural representation regions for both types of memory, were observed in the frontal/parietal cortices, the limbic system (including HPC), and the cerebellum. Abbreviations: L, left hemisphere; R, right hemisphere; Ant, anterior; Post, posterior; ACG, anterior cingulate; AG, angular gyrus; CAU, caudate; FP, frontal pole; HPC, hippocampus; HYPO, hypothalamus; IFG, inferior frontal gyrus; INS, insular cortex; ITG, inferior temporal gyrus; LG, lingual gyrus; LOC, lateral occipital cortex; vmPFC, ventromedial prefrontal cortex; MFG, middle frontal gyrus; MTG, middle temporal gyrus; OFA, occipital face area; OFC, orbitofrontal cortex; PaCiG, paracingulate gyrus; PCG, posterior cingulate; PCUN, precuneous cortex; PrimV, primary visual areas; SC, superior colliculus; SFG, superior frontal gyrus; SMA, supplementary motor cortex; SMG, supramarginal gyrus; SubCalC, subcallosal cortex; TOFusc, temporal occipital fusiform cortex; TP, temporal pole.

#### 2.3.2 Widespread neural representations of remote memories in the frontal-parietal cortices and the limbic system

Whole-brain searchlight analysis revealed neural representations of remote memories in 36 brain regions, with 22 of these located in the frontal-parietal cortices. Specifically, the frontal regions included the bilateral regions in the frontal pole (FP), inferior frontal gyrus (IFG), and superior frontal gyrus (SFG); the left hemisphere regions of the middle frontal gyrus (MFG), paracingulate (PaCiG), the subcallosal cortex (SubCalC), and the insula (INS); as well as the right OFC. The parietal regions included the bilateral supramarginal gyrus (SMG) and PCUN; the left postcentral gyrus (POCG.L); and the right angular gyrus (AG.R). Additionally, remote memories were found in six limbic areas, including the bilateral dorsal anterior cingulate gyrus (dACG), the right posterior cingulate (PCG.R), the aHPC.L, the pHPC.R, the left anterior parahippocampal gyrus (aPHG.L), and the bilateral hypothalami (HYPO). These findings indicate that the majority of neural representations of remote memories are distributed across the frontal-parietal cortices and the limbic system.

Furthermore, neural representations of remote memories were identified in occipital, temporal, and cerebellar regions. These include four areas in the left lingual gyrus (LG.L), the right inferior lateral occipital cortex (iLOC.R), and the bilateral superior LOC (sLOC); one area in the temporal region, the right temporal pole (TP.R), and three areas in the cerebellum, namely the right cerebellum 6 (Cereb6.R), the left Crus2 (Cereb2.L), and the left cerebellum 7b (Cereb7.L).

#### 2.3.3 Widespread neural representations of recent memories in the frontal and occipital cortices

Neural representations of recent associative memories were distributed across 55 brain regions (Fig. 2b and Supplementary Table 3). They were most prominent in the frontal cortex, encompassing 19 regions in both the medial and lateral areas, including the left vmPFC (vmPFC.L), the bilateral SFG, PaCiG, and ACG on the medial side, as well as the bilateral FP, MFG, and precentral gyrus, as well as the right IFG and the right OFC on the lateral side. The occipital cortex also showed extensive neural representations, notably in 10 regions of the ventral visual pathway from the right primary visual cortex V1 to the secondary visual cortices V2, including the occipital pole (OP), iLOC, occipital fusiform gyrus (OFusG) in both hemispheres, the bilateral occipito-temporal regions (TOFusC), and finally reached the left inferior temporal gryus (ITG.L).

Neural representations of recent memories were also widespread in eight cerebellar regions involved in motor processing and task-related attentional/executive processing, including the left cerebellum 8 (Cereb8.L), the left Crus1 (Cereb1.L), the Cereb6.R, the right Crus2 (Cereb2.R), and the right cerebellum 3 (Cereb3.R) (Supplementary Figure 2b).

Furthermore, neural representations of recent memories were observed in the left SMG and bilateral precuneus (PCUN) in the parietal cortex; the TP.R and the right posterior middle temporal gyrus (pMTG.R) in the temporal cortex; and the pHPC.R, bilateral ACG, and the left caudate (CAU.L) in the limbic system.

#### 2.3.4 Distinct neural representation regions for remote and recent memories

Our cross-modal MVPA findings highlighted distinct neural representations for remote and recent memories throughout the brain, with remote memories showing broader distribution than recent memories in the limbic system and parietal cortex, whereas recent memories were more widespread in the frontal, occipital cortices, and the cerebellum.

Upon analyzing their overlap regions, we discovered several areas in the frontal cortex (OFC.R, SFG.R, FP.L, and SubCalC), parietal cortex (SMG and PCUN), cerebellum (Cereb6.R), and limbic system (ACG and pHPC.R) that contain neural representations of both remote and recent memories (Fig. 2c and Supplementary Table 4).

### 2.4 Enhanced global functional connectivities between the neural representation regions of remote memories

We performed gPPI analysis on the MVPA results to explore interactions between brain regions during the retrieval of remote and recent memories. In this analysis, the strength of task-dependent functional connectivity between pairs of regions under the given psychological conditions was indexed by the beta values (i.e., parameter estimates) from the gPPI model. Larger beta values indicate stronger task-dependent coupling between regions (see Methods 4.6). A one-tailed t-test with FDR correction on these beta values identified 40 significant functional connectivities (*P* < 0.05, FDR corrected) across the 36 neural representation regions of remote memories (Fig. 3a, Supplementary Table 5). However, across the 55 neural representation regions of recent memories, although 10 functional connectivities were significant at an uncorrected *P* < 0.001 , none remained significant after FDR correction (Fig. 4a, Supplementary Table 6).

**Figure 3.**
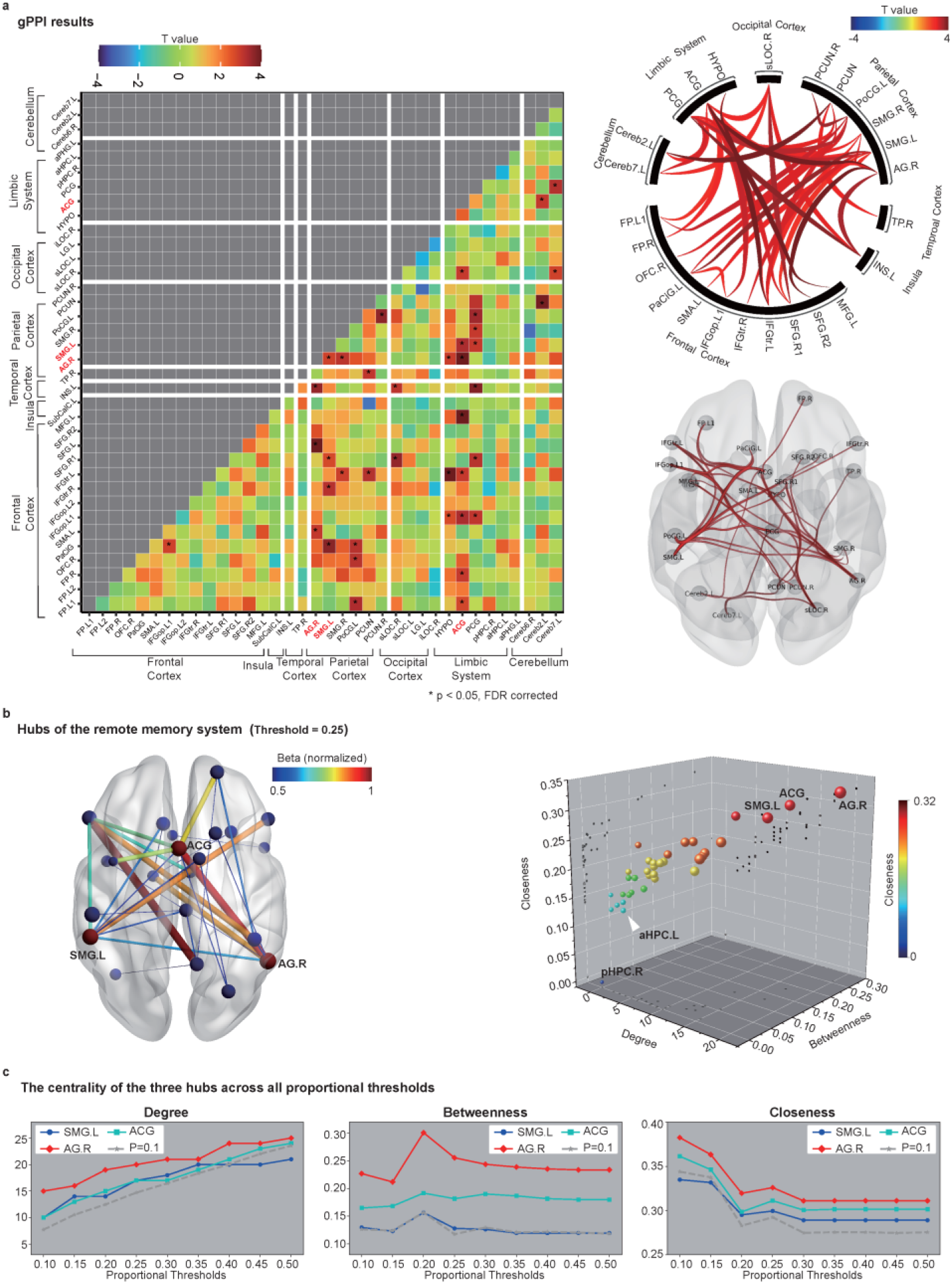
Functional connectivity and hub structure of the remote memory retrieval network. **a.** Heatmap (left) shows functional connectivities among brain regions representing remote memories. The connectogram and glass brain (right) display significant connections thresholded at *P* < 0.05 (FDR-corrected). Global rather than local connectivities were predominant (see Supplementary Table 5). **b.** Graph-theoretical analysis. Functional connectivities (gPPI beta values) were normalized to the range [0, 1]. The top 25% of connectivities (left) were used to compute degree, betweenness, and closeness centrality for each node. A three-dimensional scatterplot (right) highlights two parietal regions (AG.R, SMG.L) and one limbic region (ACG) as hubs, each showing the highest values across these metrics (*P* < 0.1 for each measure). Both panels reflect data averaged across all 30 participants. **c.** Centrality of the three identified hubs remained consistently high across proportional thresholds (PTs; 10−50% in 5% increments).

**Figure 4.**
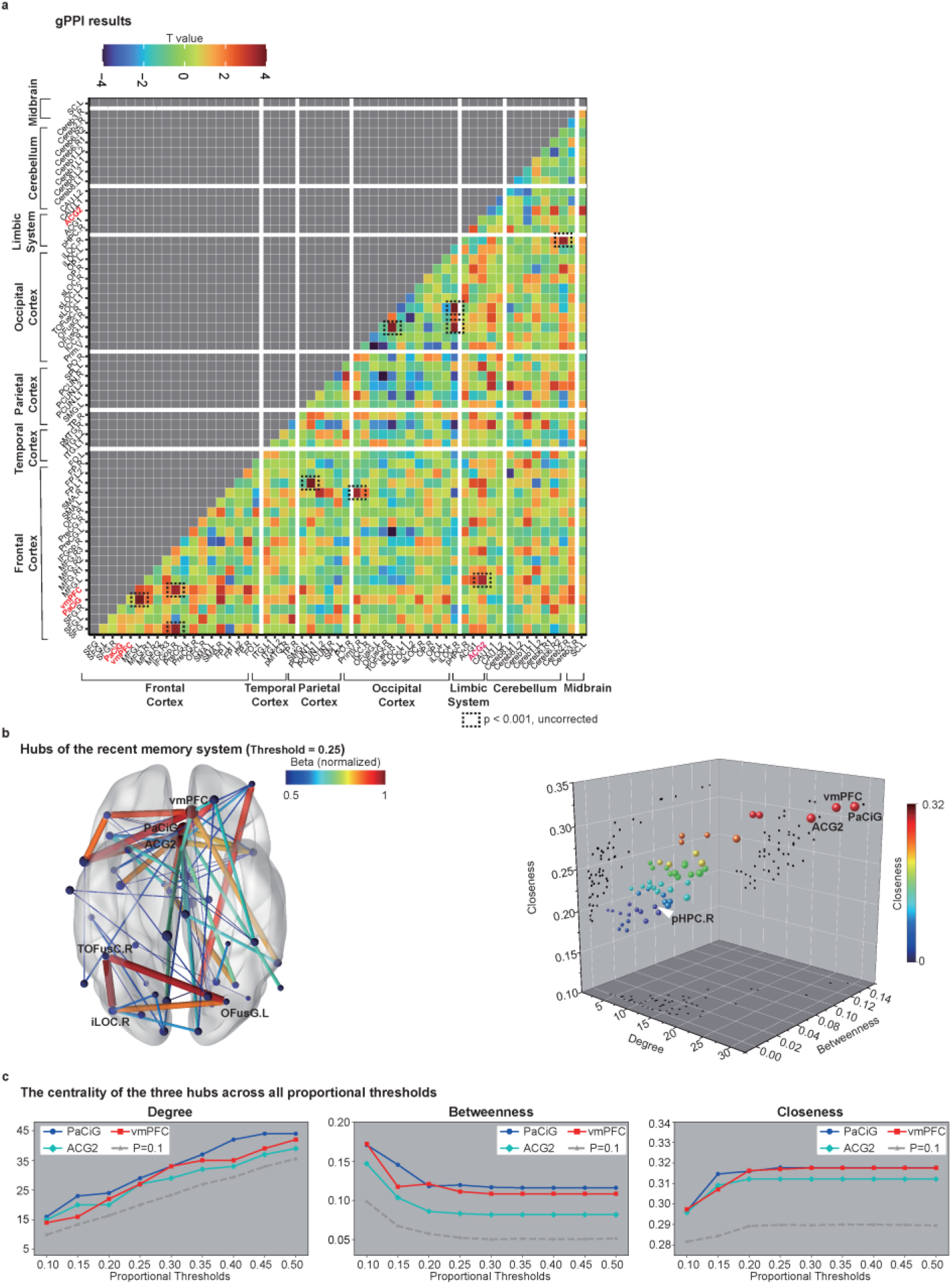
Functional connectivity and hub structure of the recent memory retrieval network. **a.** Heatmap (left) depicts functional connectivities among brain regions representing recent memories. Ten connections reached significance at an uncorrected threshold (*P* < 0.001), but none survived FDR correction (see Supplementary Table 6). **b.** Graph-theoretical analysis. The top 25% of normalized gPPI-based functional connectivities (beta values normalized to the range [0, 1]; left) were used to compute degree, betweenness, and closeness centrality for each node. Three midfrontal regions (vmPFC, PaCiG, and ACG) were identified as hubs, each showing the highest degree, betweenness, and closeness centrality (*P* < 0.1 for each metric). **c.** Centrality of these three hubs remained consistently high across proportional thresholds (PTs, 10−50% in 5% increments).

Moreover, global functional connectivities (between different brain lobes) were more pronounced than local functional connectivities (within the same lobe) in the remote memory network, with 36 being global out of 40 significant connectivities. The majority of global functional connectivities were observed between the frontal and parietal regions (11 connectivities) and between the limbic system and the frontal-parietal regions (17 connectivities).

However, local functional connectivities were more pronounced in the recent memory retrieval network. Of the 10 significant functional connectivities (*P* < 0.001, uncorrected), six were local, located in the frontal and occipital cortices.

Although the MVPA results indicated that the HPC maintains neural representations of both remote and recent memories, no significant functional connectivities involving the HPC were detected during the retrieval of either type of memory. However, by reducing the significance threshold, we identified significant functional connectivities between HIP.L and sLOC.L (beta value = 0.071 , uncorrected *p* = 0.017), as well as with PaCiG (beta value = 0.060 , uncorrected *p* = 0.011) during remote memory retrieval; significant functional connectivities were observed between HIP.R and sLOC (beta value = 0.084, uncorrected *p* = 0.008), as well as with ACG1 (beta value = 0.071, uncorrected *p* = 0.008) during recent memory retrieval.

### 2.5 The multi-hub network system in remote memory retrieval

Graph theory analyses on the memory retrieval networks identified network hubs that integrate brain functions and facilitate global neural communication during memory retrieval (see Methods 4.7) ^31, 32^. The regions with the highest degree, betweenness, and closeness centrality (*P* < 0.1 for each measure) were identified as hubs ^33^. The stability of these measures was confirmed by applying proportional thresholding (PT%: 10%–50% in 5% increments) to the functional networks (Fig. 3b).

Three hubs were identified in the remote memory retrieval network: the dACG within the mPFC, bilateral inferior parietal lobule regions AG.R and SMG.L. These regions exhibit robustly high centrality across different thresholds, with AG.R displaying the highest degree, betweenness, and closeness centrality (Fig. 3bc). For recent memory retrieval, the network’s hubs were located in the mPFC, including the vmPFC, PaCiG and ACG (Fig. 4bc). These findings suggest that, following long-term learning, memory retrieval hubs shift from being centered in the mPFC to being distributed across the frontal-parietal regions.

## 3. Discussion

One of the central aims of this study was to clarify the role of the hippocampus (HPC) in systems consolidation by identifying neural representations of both recent and remote memories. We found that recent memories were represented exclusively in the right posterior hippocampus (pHPC.R), whereas remote memories were distributed across both the pHPC.R and the left anterior hippocampus (aHPC.L). The overlap in pHPC.R suggests continuity across memory age, while the additional recruitment of aHPC.L for remote memories aligns with Trace Transformation Theory (TTT): the posterior hippocampus supports detailed recollection, whereas the anterior hippocampus contributes to gist-based retrieval ^2, 13^. These findings extend prior MVPA work decoding autobiographical memories from hippocampal activation ^21, 22^, while offering more consistent localization across participants due to our controlled face–name associative paradigm.

Our results also illuminate how memory representations shift from perceptual to semantic pathways over time. For recent memories, cross-modal MVPA and univariate analyses confirmed a strong reliance on the ventral visual stream (V1/V2 to the inferior temporal gyrus), consistent with reinstatement theories of memory that propose retrieval reactivates original sensory cortices ^34, 35^. This pathway, well established as the core system for face perception ^36, 37, 38^, was clearly engaged during recent memory retrieval but contributed minimally to remote retrieval. Remote memory signals in sensory cortices were confined to early visual regions (V1/V2), suggesting that perceptual reinstatement is time-limited.

How then are remote face–name associations retrieved? Our MVPA identified the right temporal pole (TP.R), a key node in the extended face recognition system that links perceptual inputs with person-specific semantic knowledge ^39^. In addition, the MVPA results identified a lateral visual pathway in the occipital-temporal cortex, which transmits visual information from the primary visual cortex (V1) through the posterior superior temporal sulcus (pSTS) to TP.R ^37, 40^. This lateral visual pathway is thought to accelerate recognition of familiar faces through semantic integration ^41^. Together, these findings support a representational shift: recent memories rely on perceptual reinstatement, whereas remote memories are mediated by semantic pathways and extended face recognition regions.

Beyond the TP.R, we observed remote memory representations across a broad set of extended network regions, including the insula, anterior hippocampus, posterior cingulate gyrus, amygdala, orbitofrontal cortex, hypothalamus, and retrosplenial/ACC. Compared to recent memories, this distribution was more widespread and weighted toward regions linked to semantic processing and emotion. This expansion suggests a gradual semanticization of memory content ^15,42,43^, consistent with behavioral evidence that remote memories contain fewer perceptual details but richer emotional and personal significance.

The cerebellum also mirrored this transformation. While both recent and remote memories were represented bilaterally, recent memories were more prominent in the left cerebellum (linked to picture-based recall), whereas remote memories localized to non-motor cerebellar regions associated with attention and executive control ^44, 45^. This further supports the idea that recent retrieval depends more on perceptual traces, whereas remote retrieval depends more on semantic integration.

Neural representations of memory are reflected not only in activation patterns, but also in functional connectivities ^43, 46^. Remote memory retrieval was associated with globally enhanced neocortical and limbic connectivity, reflecting a stabilized, integrated network. By contrast, recent memory retrieval showed weaker and less stable connectivity, coupled with stronger frontal activations (IFG, SFG, OFC) consistent with the need for greater cognitive control to support fragile associations ^47^. Increased PCC/ PCUN engagement during recent retrieval also overlapped with regions preferentially engaged by episodic (vs. semantic) recall ^47^, underscoring that recent memories remain episodic in nature, while remote memories have undergone semantic transformation.

A striking finding was the reorganization of network hubs. While TTT emphasizes the vmPFC as the primary schema hub during consolidation ^14^, we observed a shift over time: from vmPFC dominance during recent retrieval to a distributed hub structure for remote memories, centered on the medial PFC and bilateral inferior parietal lobule (SMG.L, AG.R). Notably, AG.R showed the highest centrality, consistent with its role as a semantic and conceptual knowledge hub ^14, 48, 49^. This transition echoes Sommer’s long-term training results ^15^, where vmPFC activity declined while semantic regions gained prominence. We propose that with long-term repetition, episodic retrieval is increasingly scaffolded by semantic frameworks, with parietal regions integrating conceptual knowledge into retrieval processes ^29, 50, 51^.

Synthesizing these findings, we propose a two-phase model of systems consolidation (Fig. 5). Phase 1: Schema-mediated consolidation. Recently formed memories rely on perceptual reinstatement within low-level visual regions and unstable functional connectivities. Retrieval requires strong cognitive control, with vmPFC acting as a schema hub to integrate new episodic traces into existing knowledge networks (Fig. 5, left) ^15, 52, 53^.

**Figure 5.**
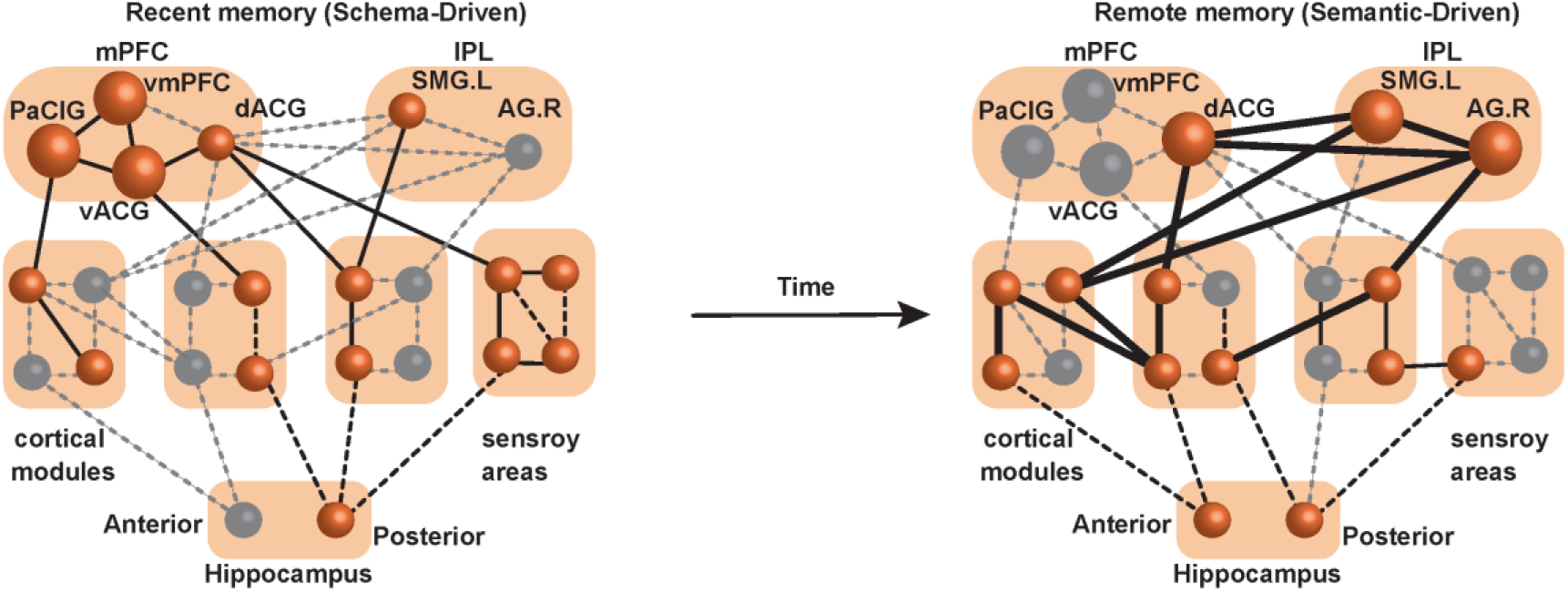
Two-phase memory trace transformation model. **1. Initial phase.** Memory retrieval relies on processes resembling those engaged during encoding, including reactivation of primary sensory regions. Functional connectivities among memory-representation regions are not yet stabilized, and retrieval requires high levels of cognitive control. In this stage, the vmPFC acts as the central hub of the recent memory retrieval network. **2. Post-initial phase.** With ongoing systems consolidation, the memory network expands to incorporate a broader set of semantic memory regions. Functional connectivities become globally stable, and retrieval is less dependent on the vmPFC, instead drawing on multiple hubs distributed across the frontal and parietal cortices. At this stage, the remote memory retrieval network emerges as an integrated system characterized by robust and enduring global functional connectivities among relevant brain regions.

Phase 2: Semantic-mediated consolidation. With long-term repetition, memory representations shift toward high-level semantic regions (TP.R, AG.R), supported by the lateral visual pathway. Functional connectivity becomes globally stabilized, and hub roles transfer from vmPFC to mPFC and inferior parietal lobule. Retrieval in this phase is mediated by semantic frameworks rather than perceptual reinstatement (Fig. 5, right).

This model explains the process of systems consolidation in long-term repetitive learning and effectively explains memory deficits observed in neurodegenerative disorders, such as Alzheimer’s disease (AD) and semantic dementia (SD) ^29^. During Phase 1, memory retrieval relies on the ventral visual pathway; damage to the medial temporal lobe can disrupt visual information processing during memory retrieval, leading to difficulties in remembering sensory-perceptual details in individuals with AD ^29^. In contrast, during Phase 2, remote memories have shifted to high-level semantic representation regions. The retrieval of remote memories relies on the lateral visual pathway, independent of the medial temporal lobe, potentially explaining why remote memories are less impaired in AD patients. Conversely, individuals with SD suffer from degeneration in the lateral temporal lobe, making their remote memories more susceptible to impairment ^54, 55^.

Although remote memories predominantly rely on semantic regions, they still retain representations within the hippocampus (HPC). Interestingly, we observed no significant hippocampal connectivity with other regions during retrieval. Previous studies suggest that such connectivity is highly task-dependent ^34^. For example, McCormick et al. reported that the anterior hippocampus (aHPC) exhibited connectivity with the medial prefrontal cortex (mPFC) during memory construction, whereas the posterior hippocampus (pHPC) showed connectivity with the posterior neocortex during memory elaboration ^56^. In contrast, Wang et al. found no significant functional connectivity between the HPC and other brain regions when participants retrieved information about a person’s social attributes, such as occupational status ^57^. Accordingly, our finding of no significant hippocampal connectivity during either recent or remote memory retrieval may reflect the specific demands of our experimental paradigm.

Our study leveraged an innovative paradigm using Pokémon face–name associations, which uniquely allowed for shared recent and remote memories across participants. This approach minimized variability inherent to autobiographical tasks while capturing long-term, ecologically valid memory representations shaped by repetitive childhood learning. Nevertheless, our design compared only two time points: one day (recent) and over a decade (remote). Future work should adopt longitudinal designs spanning multiple retention intervals to map consolidation dynamics more continuously.

Together, these findings refine existing theories of systems consolidation. We demonstrate that the hippocampus retains long-lasting representational roles while the broader network undergoes large-scale reorganization—from perceptual reinstatement and vmPFC schema integration to semantic hubs in parietal cortices. By proposing a two-phase model of memory transformation, we provide a framework that unifies prior discrepancies and links basic consolidation processes to clinical memory disorders.

## 4. Methods

### 4.1 Participants

A total of 32 healthy university students (six females; age: 19.8 ± 1.7 (mean ± SD) yr, range: 18–24 yr) with normal or corrected-to-normal vision participated in the experiments. Participants were recruited based on the following criteria: previous extensive experience of playing Pokémon Diamond/Pearl (released in 2006) at the time of its release, as well as little or no experience with the latest Pokémon Sword & Shield (released in 2019) at the time of the experiment (2020). Two of the participants were excluded from data analyses due to excessive head movements (mean framewise displacement > 0.25 mm) or falling asleep during MRI scanning. This study was approved by the Ethics Committee of Kochi University of Technology, and all participants provided written informed consent prior to participation.

### 4.2 Procedures

The experiments were conducted in two consecutive days (Fig. 1b).

On Day 1, participants were required to play the 2019-Pokémon game on a Nintendo Switch Lite, controlling three major characters: Grookey, Scorbunny, and Sobble. Participants controlled each character for 25 mins without any other instruction. It was expected that participants would incidentally learn the face-name associations of these characters during the gameplay.

On Day 2, participants underwent fMRI scanning while completing a face-name association task (Fig. 1b). This task involved the retrieval of both remote and recent memories. The recent memory items consisted of the faces and names of three 2019-Pokémon characters that participants had learned the previous day, while the remote memory items consisted of faces and names of three major 2006-Pokémon characters (Piplup, Turtwig, and Chimchar). During the face-name association task, participants were shown a Pokémon character’s face or name (in Japanese Katakata characters) and required to identify the corresponding name or face. The stimuli were presented on a BOLDscreen (Cambridge Research Systems Ltd., UK) with a visual angle of 8◦ × 6◦ against a grey background (RGB: [160, 160, 160]). All stimuli were calibrated for equal luminance and contrast using the RGB-shine toolbox in MATLAB ^46^.

The face-name association task was conducted using PsychToolbox-3 in MATLAB R2019b (Math-Works, Natick, MA) and consisted of four runs. Each run started with a 20-second fixation cross, followed by a series of 36 trials. In each trial, a Poké ball was presented on the center of the screen. To start the trial, participants were asked to press a right-handed button as soon as possible. A character’s name or face was then presented for 2 s. After a delay interval (11 s, or 13 s), the experiment continued with the next trial. The total duration of one run was approximately 10 mins.

Following the task, participants’ face-name association memories were evaluated by a surprise name-writing test. The faces of the Pokémon characters used in this study were printed on a sheet of paper, and participants were asked to write down the corresponding names.

### 4.3 fMRI data acquisition and preprocessing

While participants were performing the face-name association task, whole-brain scanning was conducted on a Siemens Magnetom Prisma 3T MR scanner with a 64-channel head coil. A high-resolution T1-weighted anatomical image was acquired using a MPRAGE sequence (repetition time [TR] = 1900 ms; echo time [TE] = 2.52 ms; flip angle [FA] = 9^°^; field of view [FOV] = 250 mm; matrix = 256 × 256; in-plane resolution = 1 × 1 mm^2^, slice thickness = 1 mm; 176 slices). Functional images were collected using a multiband echo planar imaging (EPI) pulse sequence (TR = 743 ms; TE = 35.6 ms; FA = 48^°^ ; FOV = 192 mm; matrix = 96 × 96 ; in-plane resolution = 2 × 2 mm^2^, slice thickness = 2 mm; 72 slices; Multi-band acceleration factor = 8). fMRIPrep (version 22.0.1, https://fmriprep.org/en/stable/) was used for preprocessing anatomical and functional data ^58^.

#### 4.3.1 Anatomical data preprocessing

The T1-weighted (T1w) image was corrected for intensity non-uniformity using N4BiasFieldCorrection ^59^ in ANTs 2.3.3 ^60^ and used as the T1w-reference. ANTs was then used to skull-strip the T1w-reference, resulting in the brain-extracted T1w. Brain tissue segmentation of cerebrospinal fluid (CSF), white-matter (WM), and gray-matter (GM) was carried out on the brain-extracted T1w using fast in FSL 6.0.5.1 ^61^. Brain surface reconstruction was accomplished by recon-all in FreeSurfer 7.2.0 ^62^. The T1w image was then spatially normalized to a standard space, the ICBM 152 Nonlinear Asymmetrical template version 2009c (MNI152NLin2009cAsym) ^63^.

#### 4.3.2 Functional data preprocessing

A BOLD reference volume was generated and co-registered to the T1w reference. The BOLD timeseries were resampled into their original, native space. Several confounds, framewise displacement (FD), derivative of root mean square variance over voxels (DVARS), CSF, WM, and the whole-brain global signals were estimated from the preprocessed BOLD time-series. Furthermore, a set of physiological regressors were extracted to enable component-based noise correction. The BOLD time-series were then resampled into the standard space (MNI152NLin2009cAsym).

To obtain unsmoothed BOLD signals required for subsequent analyses, noises identified by independent component analysis-based automatic removal of motion artifacts (ICA-AROMA) ^64^ were removed from the BOLD signals using FSL’s MELODIC function instead of the default denoising provided by fMRIprep.

### 4.4 Univariate General Linear Model Analyses (GLM)

A GLM analysis was carried out using FLS with task regressors of recent memory (2019-Pokémon characters’ names and faces) and remote memory (2006-Pokémon characters’ names and faces). To reduce noise effects, two additional noise confounds, the global signal and outlier scans, were also included. Before conducting the GLM analysis, the denoised fMRI data were smoothed using a Gaussian kernel (FWHM = 6 mm) to increase sensitivity.

For group analysis, a one-sample t-test with 10,000 permutations was used to compare the recent to remote memory (“Recent − Remote”) and remote to recent (“Remote − Recent”) across all 30 participants. The FDR multiple correction of the activation maps was performed within a gray matter mask (mean probability > 10%) using LIPSIA 3.1.0 (https://github.com/lipsia-fmri/lipsia). The LIPSIA adopted a non-parametric and threshold-free framework (LISA) ^65^ for group-level analysis, which has been shown to be a sensitive and robust method and has been validated in previous GLM and MVPA studies ^66, 67^.

### 4.5 Cross-modal decoding MVPA

In the event-related MVPA, beta maps are employed as observation vectors to enhance identification accuracy. The Least Squares-Separate (LSS) model ^68^ was utilized to calculate the beta maps for each event from the denoised fMRI data using Nistats NiBetaSeries v0.6.0 (https://nibetaseries.readthedocs.io/en/stable/) ^69^. The global signal and outlier scans were included as nuisance regressors in the LSS model. Note that, in contrast to the GLM analysis, no spatial smoothing was applied here.

Cross-modal MVPA implemented by BrainIAK (https://brainiak.org/) ^70^ was employed. To investigate neural representations of remote associative memories, we first trained a linear support vector classifier (SVC, cost parameter = 1, maximum number of iterations = 5,000) using beta maps of the names of the three 2006-Pokémon characters and then tested it on the beta maps of the faces of these characters. Conversely, the classifier was trained on data from face stimuli and tested on data from name stimuli. In this way, we calculated the average cross-modal (face-name) decoding accuracy of remote memories in a searchlight strategy across all voxels of the brain (search radius = 9 mm). The outline of the cross-modal decoding strategy is illustrated in Fig. 1c.

Above-chance accuracy maps were obtained by subtracting the chance-level (33.4%) from the decoding accuracy maps. A one-sample t-test with 10,000 permutations was performed on the above-chance accuracy maps, followed by FDR multiple correction to identify significant positive voxels (*P* < 0.01). This process was accomplished by LIPSIA, and the significant result was constrained to the gray matter by a gray matter mask (mean probability > 10%).

The same analysis was applied to the fMRI data from the 2019-Pokémon characters to investigate neural representations of recent memories.

### 4.6 Task-based functional connectivity with gPPI

Generalized psychophysiological interaction (gPPI) ^71^ implemented in the CONN toolbox was used to explore the functional connectivities between memory-related regions obtained by the cross-modal decoding MVPA.

#### 4.6.1 Pre-processing

The fMRI data that had been preprocessed using fMRIPrep were imported into the CONN toolbox for further processing. Linear regression was applied to remove potential confounding effects from the data, which were subsequently high-pass filtered (> 0.008 Hz). The CONN’s default settings for potential confounding variables included anatomical noise, motion artifacts arising from participants’ head movement, and constant task effects ^72^. The anatomical noise was composed of five principal components of WM and five principal components of CSF, which were calculated by anatomical CompCor (aCompCor) ^73^. The participants’ head-motion realignment consisted of six head-motion parameters (three rotations and three translations) and their first-order temporal derivatives, as well as time frames with excessive motion ^74^. These anatomical and motion-related confounds were percalculated with fMRIPrep. The constant task effects were composed of standard task responses (i.e., task regressors convolved with a canonical two-gamma hemodynamic response function) and their first-order derivatives.

#### 4.6.2 ROI-to-ROI gPPI analyses in CONN

In the ROI-to-ROI gPPI analyses, it is important to carefully consider the use of spatial smoothing, as it can potentially lead to the incorporation of unrelated voxels, thus altering the characteristics of the functional networks, such as the degrees of nodes ^75^. To avoid this issue, no spatial smoothing was applied here. Instead, the averaged BOLD signal of each ROI was used as a physiological regressor. A separate multiple regression model was executed for each pair of ROIs. The multiple regression of a target ROI signal *R_j_*(*t*) on a seed ROI signal *R_i_*(*t*) is described by the following equation:

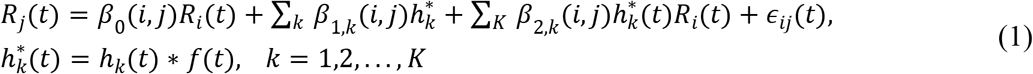

where the *k*-th psychological regressor, ℎ^∗^ (*t*), is obtained by convolving the task onset information ℎ*_k_*(*t*) with a canonical hemodynamic response function *f*(*t*) . The optimal values of *β*_0_(*i*, *j*) , *β*_1,*k*_(*i*, *j*), and *β*_2,*k*_(*i*, *j*) can be estimated by minimizing the residuals *ε_ij_*(*t*) using an Ordinary Least Squares (OLS) solution. This process is represented by the following equation:]

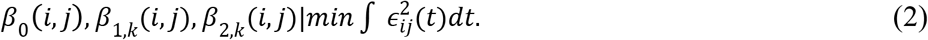

The parameter *β*_2,*k*_(*i*, *j*) represents directional effective connectivity from the seed region *R_i_* to the target region *R_j_* under the *k*-th psychological condition ℎ^∗^ (*t*). In contrast, *β*_2,*k*_(*j*, *i*) represents directional effective connectivity from *R_j_* to *R_i_*. Our focus was on the functional connectivity between regions, rather than on directional effective connectivity. Therefore, the strength of functional connectivity between regions *R_i_* and *R_j_* was defined as the average of *β*_2,*k*_(*i*, *j*) and *β*_2,*k*_(*j*, *i*).

#### 4.6.3 gPPI analyses of remote and recent memory

According to the results of the cross-modal decoding analysis, it was discovered that the neural representations of recent memories related to the 2019-Pokémon characters were present in 55 brain regions, whereas the neural representations of remote memories related to the 2006-Pokémon characters were present in 36 brain regions. To investigate the interactions among various brain regions in facilitating the retrieval of remote and recent memories, gPPI analyses were conducted separately for remote and recent memories in their respective relevant regions. In addition, significant positive functional connectivities between brain regions were found by employing a one-tailed t-test with FDR multiple comparison corrections in R (version 4.1.2).

### 4.7 Graph theory analyses

Three measures degree, betweenness, and closeness centrality are commonly employed to quantify the centrality of a node within a network ^31^. These measurements were calculated by the Brain Connectivity Toolbox (BCT) ^31^ in Matlab R2021b for both the remote and recent memory networks in the following steps.

#### 4.7.1 Normalization and threshold

For each participant’s functional connectivity data, a weighted, undirected network was obtained by replacing negative connectivity values with zero and rescaling the values to the range [0, 1]. Next, the weighted connectivity network was thresholded proportionally at the top 5–50%, in increments of 5%.

#### 4.7.2 Degree

The degree of a node is defined as the number of edges connected to it. For a *n*-node network, the degree of a node *i* can be calculated by the following equation:

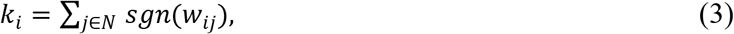

where *w_ij_* is the weighted functional connectivity between nodes *i* and *j*, *N* is the set of all nodes, and *sgn* stands for the sign function.

The “degrees_und” function from BCT was adopted to calculate degree, discarding the weight information.

#### 4.7.3 Betweenness centrality

Betweenness centrality quantifies the number of times a given node lies on the shortest path between two other nodes ^31^, as in the following equation:

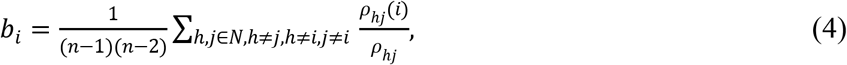

where *ρ*_ℎ*j*_ is the number of shortest paths between nodes ℎ and *j*, and *ρ*_ℎ*j*_(*i*) is the number of shortest paths between ℎ and *j* those pass-through node *i*. *n* is the total number of nodes.

The weighted connectivity networks were first converted into length matrices using the “weight_conversion” function. These matrices were further processed by the “betweenness_wei” to calculate the between-ness centralities of network nodes.

#### 4.7.4 Closeness centrality

Closeness centrality is the reciprocal of the average shortest path length between a node to all other nodes, as defined below.

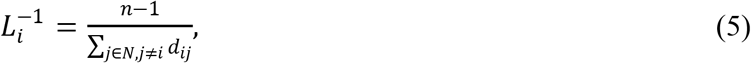

where *d_ij_* is the shortest paths between nodes *i* and *j*, and *n* is the total number of nodes. Here, the shortest paths between nodes were calculated by function “distance_wei”.

#### 4.7.5 Hub analysis

Nodes with highest degree, betweenness, and closeness centrality (for each measure *P* < 0.1) can be identified as brain hubs, which is in line with the study by Agler et al. ^33^.

The data distributions of degree and closeness centrality in our results were confirmed to follow a normal distribution, whereas the data of betweenness centrality exhibited a log-normal distribution. To establish the threshold for *P* < 0.1, bootstrap samples were generated for each measurement by parametric bootstrap with 1000 iterations, and the threshold was set at the 90% percentile point (quantile) of the resulting bootstrap samples.

## Author contributions

R.W. and K.N. conceived the project. R.W., R.K., S.M. and K.N. collected data. J.F., R.W., R.K. and K.N. analyzed data. R.W., I.H., K.J. and K.N. wrote the manuscript.

## Acknowledgement

We would like to thank Ms. Maoko Yamanaka for her administrative assistance. This study was supported by KAKENHI from Japan Society for the Promotion of Science (23H00413 to K.N. and I.H), and by AMED under grant number JP26wm0625205 to K.N. and I.H.

## References

[1] Squire, L. R. & Alvarez, P. Retrograde amnesia and memory consolidation: a neurobiological perspective. Curr. Opin. Neurobiol. 5, 169–177 (1995).

[2] Sekeres, M. J., Winocur, G. & Moscovitch, M. The hippocampus and related neocortical structures in memory transformation. Neurosci. Lett. 680, 39–53 (2018).

[3] Zola-Morgan, S. M. & Squire, L. R. The primate hippocampal formation: evidence for a time-limited role in memory storage. Science 250, 288–290 (1990).

[4] Maviel, T., Durkin, T. P., Menzaghi, F. & Bontempi, B. Sites of neocortical reorganization critical for remote spatial memory. Science 305, 96–99 (2004).

[5] Scoville, W. B. & Milner, B. Loss of recent memory after bilateral hippocampal lesions. J. Neurol. Neurosurg. Psychiat. 20, 11–21 (1957).

[6] Cipolotti, L., et al. Long-term retrograde amnesia. . . the crucial role of the hippocampus. Neuropsychologia 39, 151–172 (2001).

[7] Marr, D., Willshaw, D. & McNaughton, B. Simple memory: a theory for archicortex. In Vaina, L. (ed.) From the Retina to the Neocortex, 59–128 (Birkhäuser Boston, Boston, MA, 1991).

[8] Nadel, L. & Moscovitch, M. Memory consolidation, retrograde amnesia and the hippocampal complex. Curr. Opin. Neurobiol. 7, 217–227 (1997).

[9] Nadel, L., Samsonovich, A., Ryan, L. & Moscovitch, M. Multiple trace theory of human memory: Computational, neuroimaging, and neuropsychological results. Hippocampus 10, 352–368 (2000).

[10] Battaglia, F. P., Borensztajn, G. & Bod, R. Structured cognition and neural systems: From rats to language. Neurosci. Biobehav. Rev. 36, 1626–1639 (2012).

[11] McClelland, J. L., McNaughton, B. L. & O’Reilly, R. C. Why there are complementary learning systems in the hippocampus and neocortex: Insights from the successes and failures of connectionist models of learning and memory. Psychol. Rev. 102, 419–457 (1995).

[12] Sekeres, M. J. et al. Changes in patterns of neural activity underlie a time-dependent transformation of memory in rats and humans. Hippocampus 28, 745–764 (2018).

[13] Winocur, G., Moscovitch, M. & Bontempi, B. Memory formation and long-term retention in humans and animals: Convergence towards a transformation account of hippocampal–neocortical interactions. Neuropsychologia 48, 2339–2356 (2010).

[14] Gilboa, A. & Marlatte, H. Neurobiology of schemas and schema-mediated memory. Trends Cogn. Sci. 21, 618–631 (2017).

[15] Sommer, T. The emergence of knowledge and how it supports the memory for novel related information. Cereb. Cortex 27, 1906–1921 (2017).

[16] Haist, F., Gore, J. B. & Mao, H. Consolidation of human memory over decades revealed by functional magnetic resonance imaging. Nat. Neurosci. 4, 1139–1145 (2001).

[17] Watanabe, T. et al. Functional dissociation between anterior and posterior temporal cortical regions during retrieval of remote memory. J. Neurosci. 32, 9659–9670 (2012).

[18] Gilmore, A. W. et al. Evidence supporting a time-limited hippocampal role in retrieving autobiographical memories. Proc. Natl. Acad. Sci. U. S. A. 118, e2023069118 (2021).

[19] Steinvorth, S., Corkin, S. & Halgren, E. Ecphory of autobiographical memories: an fMRI study of recent and remote memory retrieval. Neuroimage 30, 285–298 (2006).

[20] Bernard, F. A. et al. The hippocampal region is involved in successful recognition of both remote and recent famous faces. Neuroimage 22, 1704–1714 (2004).

[21] Bonnici, H. M. et al. Detecting representations of recent and remote autobiographical memories in vmPFC and hippocampus. J. Neurosci. 32, 16982–16991 (2012).

[22] Bonnici, H. M. & Maguire, E. A. Two years later–Revisiting autobiographical memory representations in vmPFC and hippocampus. Neuropsychologia 110, 159–169 (2018).

[23] Chadwick, M. J., Bonnici, H. M. & Maguire, E. A. Decoding information in the human hippocampus: A user’s guide. Neuropsychologia 50, 3107–3121 (2012).

[24] Norman, K. A., Polyn, S. M., Detre, G. J. & Haxby, J. V. Beyond mind-reading: multi-voxel pattern analysis of fMRI data. Trends Cogn. Sci. 10, 424–430 (2006).

[25] Xue, G. et al. Greater neural pattern similarity across repetitions is associated with better memory. Science 330, 97–101 (2010).

[26] Rissman, J., Greely, H. T. & Wagner, A. D. Detecting individual memories through the neural decoding of memory states and past experience. Proc. Natl. Acad. Sci. U. S. A. 107, 9849–9854 (2010).

[27] Du, X. et al. Differential activation of the medial temporal lobe during item and associative memory across time. Neuropsychologia 135, 107252 (2019).

[28] Gilboa, A. & Moscovitch, M. No consolidation without representation: Correspondence between neural and psychological representations in recent and remote memory. Neuron 109, 2239–2255 (2021).

[29] Irish, M. & Piguet, O. The pivotal role of semantic memory in remembering the past and imagining the future. Front. Behav. Neurosci. 7, 27 (2013).

[30] Gomez, J., Barnett, M. & Grill-Spector, K. Extensive childhood experience with Pokémon suggests eccentricity drives organization of visual cortex. Nat. Hum. Behav. 3, 611–624 (2019).

[31] Rubinov, M. & Sporns, O. Complex network measures of brain connectivity: Uses and interpretations. Neuroimage 52, 1059–1069 (2010).

[32] van den Heuvel, M. P. & Sporns, O. Network hubs in the human brain. Trends Cogn. Sci. 17, 683–696 (2013).

[33] Agler, M. T. et al. Microbial hub taxa link host and abiotic factors to plant microbiome variation. PLoS Biol. 14, e1002352 (2016).

[34] Moscovitch, M. & Gilboa, A. Systems consolidation, transformation and reorganization: Multiple trace theory, trace transformation theory and their competitors. In The Oxford Handbook of Human Memory, Two Volume Pack: Foundations and Applications (Oxford University Press, 2024).

[35] Larzabal, C., Bacon-Macé, N., Muratot, S. & Thorpe, S. J. Tracking your mind’s eye during recollection: Decoding the long-term recall of short audiovisual clips. J. Cogn. Neurosci. 32, 50–64 (2020).

[36] Rapcsak, S. Z. Face recognition. Curr. Neurol. Neurosci. Rep. 19, 1–9 (2019).

[37] Finzi, D. et al. Differential spatial computations in ventral and lateral face-selective regions are scaffolded by structural connections. Nat. Commun. 12, 2278 (2021).

[38] She, L., Benna, M. K., Shi, Y., Fusi, S. & Tsao, D. Y. Temporal multiplexing of perception and memory codes in IT cortex. Nature 629, 861–868 (2024).

[39] Collins, J. A. & Olson, I. R. Beyond the FFA: The role of the ventral anterior temporal lobes in face processing. Neuropsychologia 61, 65–79 (2014).

[40] Pitcher, D. & Ungerleider, L. G. Evidence for a third visual pathway specialized for social perception. Trends Cogn. Sci. 25, 100–110 (2021).

[41] Landi, S. M., Viswanathan, P., Serene, S. & Freiwald, W. A. A fast link between face perception and memory in the temporal pole. Science 373, 581–585 (2021).

[42] Winocur, G. & Moscovitch, M. Memory transformation and systems consolidation. J. Int. Neuropsychol. Soc. 17, 766–780 (2011).

[43] Xue, G. From remembering to reconstruction: The transformative neural representation of episodic memory. Prog. Neurobiol. 102351 (2022).

[44] Fliessbach, K., Trautner, P., Quesada, C. M., Elger, C. E. & Weber, B. Cerebellar contributions to episodic memory encoding as revealed by fMRI. Neuroimage 35, 1330–1337 (2007).

[45] Otten, L. J. & Rugg, M. D. Task-dependency of the neural correlates of episodic encoding as measured by fMRI. Cereb. Cortex 11, 1150–1160 (2001).

[46] Keerativittayayut, R., Aoki, R., Sarabi, M. T., Jimura, K. & Nakahara, K. Large-scale network integration in the human brain tracks temporal fluctuations in memory encoding performance. Elife 7, e32696 (2018).

[47] Vatansever, D., Smallwood, J. & Jefferies, E. Varying demands for cognitive control reveals shared neural processes supporting semantic and episodic memory retrieval. Nat. Commun. 12, 2134 (2021).

[48] Binder, J. R., Desai, R. H., Graves, W. W. & Conant, L. L. Where is the semantic system? A critical review and meta-analysis of 120 functional neuroimaging studies. Cereb. Cortex 19, 2767–2796 (2009).

[49] Ralph, M. A. L., Jefferies, E., Patterson, K. & Rogers, T. T. The neural and computational bases of semantic cognition. Nat. Rev. Neurosci. 18, 42–55 (2017).

[50] Greenberg, D. L. & Verfaellie, M. Interdependence of episodic and semantic memory: Evidence from neuropsychology. J. Int. Neuropsychol. Soc. 16, 748–753 (2010).

[51] Irish, M., Addis, D. R., Hodges, J. R. & Piguet, O. Considering the role of semantic memory in episodic future thinking: evidence from semantic dementia. Brain 135, 2178–2191 (2012).

[52] Ghosh, V. E., Moscovitch, M., Colella, B. M. & Gilboa, A. Schema representation in patients with ventromedial PFC lesions. J. Neurosci. 34, 12057–12070 (2014).

[53] Spalding, K. N., Jones, S. H., Duff, M. C., Tranel, D. & Warren, D. E. Investigating the neural correlates of schemas: Ventromedial prefrontal cortex is necessary for normal schematic influence on memory. J. Neurosci. 35, 15746–15751 (2015).

[54] Graham, K. S. & Hodges, J. R. Differentiating the roles of the hippocampus complex and the neocortex in long-term memory storage: Evidence from the study of semantic dementia and Alzheimer’s disease. Neuropsychology 11, 77–89 (1997).

[55] Nestor, P., Graham, K. S., Bozeat, S., Simons, J. & Hodges, J. Memory consolidation and the hippocampus: further evidence from studies of autobiographical memory in semantic dementia and frontal variant frontotemporal dementia. Neuropsychologia 40, 633–654 (2002).

[56] McCormick, C., St-Laurent, M., Ty, A., Valiante, T. A. & McAndrews, M. P. Functional and effective hippocampal–neocortical connectivity during construction and elaboration of autobiographical memory retrieval. Cereb. Cortex 25, 1297–1305 (2015).

[57] Wang, Y. et al. Dynamic neural architecture for social knowledge retrieval. Proc. Natl. Acad. Sci. U. S. A. 114, E3305–E3314 (2017).

[58] Esteban, O. et al. fMRIPrep: a robust preprocessing pipeline for functional MRI. Nat. Methods 16, 111–116 (2019).

[59] Tustison, N. J. et al. N4ITK: improved N3 bias correction. IEEE Trans. Med. Imaging 29, 1310–1320 (2010).

[60] Avants, B. B., Epstein, C. L., Grossman, M. & Gee, J. C. Symmetric diffeomorphic image registration with cross-correlation: Evaluating automated labeling of elderly and neurodegenerative brain. Med. Image Anal. 12, 26–41 (2008).

[61] Zhang, Y., Brady, M. & Smith, S. Segmentation of brain MR images through a hidden Markov random field model and the expectation-maximization algorithm. IEEE Trans. Med. Imaging 20, 45–57 (2001).

[62] Dale, A. M., Fischl, B. & Sereno, M. I. Cortical surface-based analysis: I. Segmentation and surface reconstruction. Neuroimage 9, 179–194 (1999).

[63] Ciric, R. et al. TemplateFlow: FAIR-sharing of multi-scale, multi-species brain models. Nat. Methods 19, 1568–1571 (2022).

[64] Pruim, R. H. et al. ICA-AROMA: A robust ICA-based strategy for removing motion artifacts from fMRI data. Neuroimage 112, 267–277 (2015).

[65] Lohmann, G. et al. LISA improves statistical analysis for fMRI. Nat. Commun. 9, 4014 (2018).

[66] Ness, H. T. et al. Reduced hippocampal-striatal interactions during formation of durable episodic memories in aging. Cereb. Cortex 32, 2358–2372 (2022).

[67] Koelsch, S., Cheung, V. K., Jentschke, S. & Haynes, J.-D. Neocortical substrates of feelings evoked with music in the ACC, insula, and somatosensory cortex. Sci. Rep. 11, 10119 (2021).

[68] Abdulrahman, H. & Henson, R. N. Effect of trial-to-trial variability on optimal event-related fMRI design: Implications for Beta-series correlation and multi-voxel pattern analysis. NeuroImage 125, 756–766 (2016).

[69] Kent, J. D. & Herholz, P. NiBetaSeries: task related correlations in fMRI. J. Open Source Softw. 4, 1295 (2019).

[70] Kumar, M. et al. BrainIAK: The brain imaging analysis kit. Apert. Neuro 1 (2021).

[71] McLaren, D. G., Ries, M. L., Xu, G. & Johnson, S. C. A generalized form of context-dependent psychophysiological interactions (gppi): A comparison to standard approaches. Neuroimage 61, 1277–1286 (2012).

[72] Nieto-Castanon, A. Handbook of functional connectivity Magnetic Resonance Imaging methods in CONN (Hilbert Press, 2020).

[73] Behzadi, Y., Restom, K., Liau, J. & Liu, T. T. A component based noise correction method (CompCor) for BOLD and perfusion based fMRI. Neuroimage 37, 90–101 (2007).

[74] Power, J. D. et al. Methods to detect, characterize, and remove motion artifact in resting state fMRI. Neuroimage 84, 320–341 (2014).

[75] Alakörkkö, T., Saarimäki, H., Glerean, E., Saramäki, J. & Korhonen, O. Effects of spatial smoothing on functional brain networks. Eur. J. Neurosci. 46, 2471–2480 (2017).

